# From Plasmid to Pure Protein: Production and Characterization of SARS-CoV-2 PL^pro^

**DOI:** 10.1101/2025.03.09.642282

**Authors:** Anna De Falco, Rebecca Greene-Cramer, Ben A. Surina, Suren Zakian, Thomas B. Acton, Theresa A. Ramelot, Gaetano T. Montelione

**Affiliations:** Center for Biotechnology and Interdisciplinary Sciences, Rensselaer Polytechnic Institute, Troy, New York, 12180, USA; Department of Chemistry and Chemical Biology, Rensselaer Polytechnic Institute, Troy, New York, 12180, USA

## Abstract

Papain-like protease (PL^pro^) from SARS-CoV-2 is a high-priority target for COVID-19 antiviral drug development. We present protocols for PL^pro^ production in *Escherichia coli*. PL^pro^ expressed as a fusion with the *Saccharomyces cerevisiae* Smt3 protein (SUMO), is purified and obtained in its native form upon hydrolysis, with yields as high as 38 mg L^-1^. The protocol also provides isotope-enriched samples suitable for NMR studies. Protocols are also presented for PL^pro^ characterization by mass spectrometry, 1D ^19^F-NMR and 2D heteronuclear NMR, and a fluorescence-based enzyme assay.

**Highlights:**

- Production, purification, and biochemical analysis of native N- and C-termini PL^pro^
- High yields in *E. coli*, up to 38 mg L^-1^ using lysogeny broth.
- Supports labeled samples for inhibitor interaction studies.
- ^19^F NMR and fluorescence assays for inhibitor screening and IC_50_ determination.

**eTOC Blurb:** SARS-CoV-2 PL^pro^ is a key cysteine protease involved in viral replication and immune evasion, making it an important target for antiviral drug development. This study presents a detailed protocol for PL^pro^ production, purification, and biochemical analysis, achieving high yields in *E. coli*. The workflow includes fusion expression with a His-SUMO tag, isotope labeling for inhibitor studies, and assays for screening and quantifying inhibitors. This comprehensive guide facilitates large-scale production of active PL^pro^ for drug discovery and structural studies.

## Before you begin

Severe Acute Respiratory Syndrome Coronavirus-2 Papain-Like Protease (SARS-CoV-2 PL^pro^) is an essential cysteine protease involved in the SARS-CoV-2 life cycle. This protease is part of the largest non-structural protein 3 (NSP3, 1,945 residues, ∼212 kDa). The PL^pro^ domain (315 residues, ∼37 kDa) is located between the SARS unique domain (SUD/HVR) and a nucleic acid-binding domain (NAB).^1,2^ PL^pro^ serves two primary functions. First, it hydrolyzes the SARS-CoV-2 1ab polyprotein at multiple sites during viral replication.^1,3^ Additionally, it contributes to the virus’s ability to evade the immune response by removing Ubiquitin and ISG-15 from both viral and host targets.^4^ As a key target for antiviral drug development, a significant amount of active PL^pro^ is essential for studying its inhibition and identifying potential drug candidates.

This work outlines a comprehensive protocol for the production, purification, and biochemical analysis of PL^pro^ inhibition, which is similar to our other production protocol for a second SARS-CoV-2 protease, 3CL^pro^.^5^ High yields of active PL^pro^ (as high as 14 and 38 mg per L using minimal MJ9 media^6^ or standard lysogeny broth (LB),^7^ respectively) are routinely achieved using *Escherichia coli* cells. The PL^pro^ domain of NSP3 is initially expressed as a fusion with a codon-optimized N-terminal His-SUMO tag, which is subsequently cleaved by yeast Ulp1 protease (SUMO^pro^) to yield the native enzyme. The protocol also includes methods for producing ^15^N and/or ^13^C-enriched and ^19^F-Trp labeled samples, which are useful for investigating protein-inhibitor interactions. Furthermore, the study presents a ^19^F NMR-based assay for screening inhibitors and a fluorescence-based enzyme assay for determining IC_50_ values of PL^pro^ inhibitors.

This comprehensive guide aims to facilitate the production of large quantities of active SARS-CoV-2 PL^pro^, which is needed to characterize inhibitor binding to this critical antiviral drug discovery target and for structural biology studies.

### Expression Plasmids

#### His_6_-SUMO-PL^pro^ Expression Plasmid

Plasmid pGTM_CoV2_NSP3_003_SUMO (**Figure 1**) was obtained by cloning the synthetic coding sequence (CDS) for residues 746 to 1063 (PL^pro^ domain) of SARS-CoV-2 NSP3 into the pET15_ SUMO2_NESG expression vector,^8^ which includes a CDS for an N-terminal hexa-His tag upstream to a SUMO tag, and a SUMO^pro^ cleavage site. The plasmid has a pMB1 origin of replication and carries the ampicillin resistance gene. Plasmid pGTM_CoV2_NSP3_003_SUMO is available from AddGene (AddGene ID: 233739); for the remainder of this protocol, we will refer to the expressed protein construct as His_6_-SUMO-PL^pro^. pGTM_SUMO_PLpro_C111S and pGTM_SUMO_PLpro_C111A, identical plasmids with active-site mutation Cys111Ser and Cys111Ala, are also available from AddGene (AddGene IDs: 233740 and 234489, respectively).

**Figure 1:**
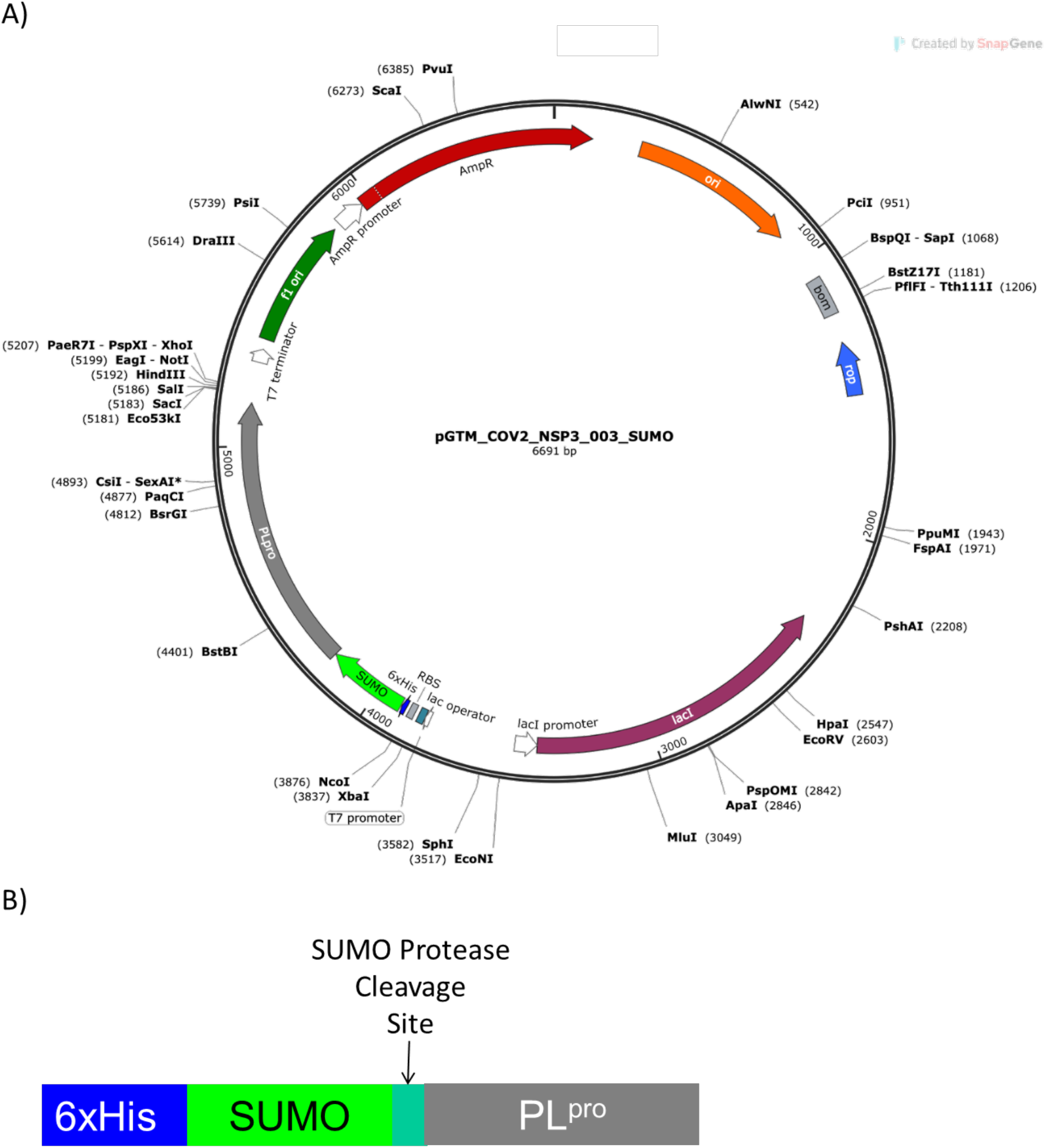
pGTM_CoV2_NSP3_003_SUMO construct design. **(A)** Final expression vector map pGTM_CoV2_NSP3_003_SUMO. Maps are generated using SnapGene® software (from Insightful Science; available at snapgene.com). The synthetic gene for PL^pro^ was obtained from Genscript, Inc., with standard Genscript codon optimization for *E. coli* expression. **(C)** Schematic representation of the expressed His_6_-SUMO-PLpro protein construct.

#### Additional Expression Plasmids

We also prepared additional constructs of PL^pro^ that have utility for biotechnology experiments (e.g., high throughput screenings). These include plasmids pGTM_SUMO-N-StrepII-PLpro (AddGene ID: 234322), pGTM_SUMO-N-8XHis-PLpro (AddGene ID: 233803) and pGTM_SUMO-C-8XHis-Plpro (AddGene ID: 234321), which provide production of His_6_-SUMO-N-StrepII-PL^pro^, His_6_-SUMO-N-His_8_-PL^pro^ and His_6_-SUMO-C-His_8_-PL^pro^, respectively. The plasmid with active-site mutation Cys111Ser for the N-terminus His tagged protein is also available from AddGene (pGTM_SUMO-N-8XHis-PLpro-C111S AddGene ID: 234472), as well as our plasmids for the production of the tryptophan protein mutants (pGTM_SUMO_PLpro_Trp93Phe, AddGene ID: 234490 and pGTM_SUMO_PLpro_Trp106Phe, AddGene ID: 234491).

### Preparation of reagent stock solutions

**Timing: 1 h**

1. Prepare 50 mg mL^-1^ ampicillin stock:
  a. Weigh 2.5 g of ampicillin sodium salt (Sigma Aldrich).
  b. Dissolve using a 50 % v/v ethanol solution to 50 mL.
  c. Sterilize using a 0.22 μm syringe filter and store at −20 °C.
2. Prepare 1M isopropyl-β-d-thiogalactopyranoside (IPTG):
  a. Weigh 2.38 g of IPTG (molar mass 238.30 g mol^-1^) (Sigma Aldrich).
  b. Dissolve in MilliQ water to a final volume of 10 mL and divide into 10 × 1 mL aliquots.
  c. Sterilize using a 0.22 μm syringe filter and store at −20 °C.
3. Prepare 1M dithiothreitol (DTT):
  a. Weigh 1.54 g of DTT (molar mass 154.25 g mol^-1^) (Sigma Aldrich).
  b. Dissolve in MilliQ water to a final volume of 10 mL and divide into 10 × 1 mL aliquots.
  c. Store at −20 °C.
4. Prepare 1 M ZnCl_2_:
  a. Weigh 6.81 g of ZnCl_2_ (molar mass 136.286 g mol^-1^).
  b. Dissolve it in a final volume of 50 mL.

**Alternative:** For the production of ^19^F-labeled-5-fluoro-L-Trp93 and Trp106 protein samples, prepare the additional stock solution:

5. Prepare 1M 5-Fluoroindole:
  a. Weigh 3.38 g of 5-Fluoroindole (molar mass 135.14 g mol^-1^).
  b. Dissolve in DMSO to a final volume of 20 mL.
  c. Store in 0.5 mL aliquots at -20 °C.

### Preparation of LB medium for bacterial growth

**Timing: 4 h**

6. Prepare 1 L Lysogeny Broth (LB) medium:
  a. Dispense 1 L MilliQ water into 2800 mL culture flask (Wilmad-LabGlass SP Scienceware).
  b. Add 20 LB Broth capsules, 1 g each (Research Products International).
  c. Sterilize in autoclave (121 °C, 15 psi, 20 min).
7. Prepare 250 mL of LB-agar for bacterial cell culture plates:
  a. Dissolve 3.75 g of agar (Sigma Aldrich) using MilliQ water to 250 mL (agar will not completely dissolve until heating during autoclave sterilization – see next points).
  b. Add 5 LB Broth capsules, 1 g each (Research Products International).
  c. Sterilize in autoclave (121 °C, 15 psi, 20 min).
  d. Allow the sterilized LB-agar mixture to cool down to ∼ 50 °C.
  e. Add 500 μL of 50 mg mL^-1^ ampicillin stock solution.

**Critical**: Antibiotics must be added after the solution has cooled to ∼ 40 °C to prevent thermal degradation.

f Under aseptic conditions, pour the LB-agar into Petri dishes.
g Allow the agar to solidify before sealing the plates with parafilm.
h Store the LB-agar plates upside down at 4 °C, to prevent aqueous vapor condensation on the surface of the LB-Agar.

### Preparation of isotope-enriched media for bacterial growth

**Timing: 1 h**

8. For ^15^N (or ^15^N,^13^C) enriched protein samples production, prepare MJ9 minimal medium calculations refer to 1 L final volume)^6^:
  a. Weigh 6 g of K_2_HPO_4_ (J.T. Baker).
  b. Weigh 9 g of KH_2_PO_4_ (J.T. Baker).
  c. Weigh 1.5 g of (^15^NH_4_)_2_SO_4_ (Sigma Aldrich).
  d. Weigh 0.5 g of sodium citrate dihydrate (J.T. Baker).
  e. Weigh between 3 and 5 g of D-Glucose or D-Glucose (U-13C6, 99 %) (Sigma Aldrich, Cambridge Isotope).
  f. Dissolve the powder in a final volume of 1 L, adding MilliQ water.
  g. Adjust the pH to 6.8.
  h. Filter using a sterile filter.
  i. Add 1 mL of vitamin stock (MEM Vitamin Solution 100× – Sigma Aldrich M6895).
  j. Add 1 g of MgSO_4_ × 7H_2_O (J.T. Baker 2500-01).
  k. Add 1 mL of trace elements stock (1000×).
  l. Add 1 mL of stock ampicillin.

### Key resources table

| REAGENT or RESOURCE | SOURCE | IDENTIFIER |
| --- | --- | --- |
| Bacterial strains |  |  |
| BL21(DE3) Competent Cells | New England Biolabs | C2527H |
| Chemicals, peptides, and recombinant proteins |  |  |
| His<sub>8</sub>-MBP-Ulp1 SUMO Protease | Prepared as described in Mazzei *et al.* 2023 |  |
| Ampicillin sodium salt | Sigma Aldrich | A9518 |
| Dithiothreitol (DTT) | Sigma Aldrich | D9779 |
| Isopropyl-β-D-thiogalactopyranoside (IPTG) | Sigma Aldrich | I6758 |
| Super Optimal broth with Catabolite repression (SOC) | Sigma Aldrich | S1797 |
| LB Broth [Lennox L Broth] 1-gram capsules | Research Products International | L24061-1000.0 |
| Agar | Sigma Aldrich | A1296 |
| Deoxyribonuclease I from bovine pancreas (DNase I) | Sigma Aldrich | DN25 |
| Nuclease Mix | Cytiva | 80-6501-42 |
| Sodium chloride (NaCl) | Sigma Aldrich | S9888 |
| Trizma base (Tris) | Sigma Aldrich | T6066 |
| Imidazole | Sigma Aldrich | I2399 |
| Zinc Chloride (ZnCl<sub>2</sub>) | Sigma Aldrich | 208086 |
| 5-Fluorindole | Gold Biotechnology | F-130-50 |
| Iron(III) chloride hexahydrate (FeCl<sub>3</sub> × 6H<sub>2</sub>O) | Sigma Aldrich | 236489 |
| Zinc sulfate heptahydrate (ZnSO<sub>4</sub> × 7H<sub>2</sub>O) | Sigma Aldrich | 221376 |
| Cobalt(II) chloride hexahydrate (CoCl<sub>2</sub> × 6H<sub>2</sub>O) | Sigma Aldrich | 255599 |
| Sodium molybdate dihydrate (Na<sub>2</sub>MoO<sub>4</sub> × 2H<sub>2</sub>O) | Sigma Aldrich | M1003 |
| Copper(II) sulfate pentahydrate (CuSO<sub>4</sub> × 5H<sub>2</sub>O) | Sigma Aldrich | 209198 |
| Boric acid (H<sub>3</sub>BO<sub>3</sub>) | Sigma Aldrich | B0394 |
| Manganese(II) sulfate (MnSO<sub>4</sub>) | Sigma Aldrich | M7506 |
| Hydrochloric acid (HCl) | Sigma Aldrich | XX0628 |
| Sodium phosphate monobasic (NaH<sub>2</sub>PO<sub>4</sub>) | Sigma Aldrich | 74092 |
| Potassium phosphate monobasic (KH<sub>2</sub>PO<sub>4</sub>) | J.T.Baker | 3246-05 |
| Potassium phosphate dibasic (K<sub>2</sub>HPO<sub>4</sub>) | J.T.Baker | 3252-01 |
| Sodium citrate dihydrate (NaC<sub>6</sub>H<sub>5</sub>O<sub>7</sub> × 2H<sub>2</sub>O) | J.T.Baker | 3646-01 |
| Magnesium sulfate heptahydrate (MgSO<sub>4</sub> × 7H<sub>2</sub>O) | J.T.Baker | 2500-01 |
| MEM Vitamin Solution (100×) | Sigma Aldrich | M6895 |
| <sup>15</sup>N-ammonium sulphate (<sup>15</sup>NH<sub>4</sub>)<sub>2</sub>SO<sub>4</sub> | Sigma Aldrich | 299286 |
| D-Glucose | Sigma Aldrich | G7520 |
| Nickel(II) sulfate (NiSO<sub>4</sub>) | Sigma Aldrich | 656895 |
| Recombinant DNA |  |  |
| pGTM_CoV2_NSP3_003_SUMO | This work – available from AddGene | AddGene ID: 233739 |
| pGTM_SUMO_PLpro_C111S | This work – available from AddGene | AddGene ID: 233740 |
| pGTM_SUMO-N-StrepII-PLpro | This work – available from AddGene | AddGene ID: 234322 |
| pGTM_SUMO-N-8XHis-PLpro | This work – available from AddGene | AddGene ID: 233803 |
| pGTM_SUMO-C-8XHis-PLpro | This work – available from AddGene | AddGene ID: 234321 |
| pGTM_SUMO-N-8XHis-PLpro-C111S | This work – available from AddGene | AddGene ID: 234472 |
| pGTM_SUMO_PLpro_Trp93Phe | This work – available from AddGene | AddGene ID: 234490 |
| pGTM_SUMO_PLpro_Trp106Phe | This work – available from AddGene | AddGene ID: 234491 |
| pGTM_YR375_SUMO_Protease_001 | Mazzei et al, 2023 | AddGene ID: 190063 |

| Other |  |  |
| --- | --- | --- |
| Sonicator 3000 | Misonix | S-3000/R2225 |
| Vivaspin 20 PES Concentrator (10,000 MWCO) (MWCO 10000 Da) | Sartorius | VS2001 |
| 5-mL HisTrap HP | Cytiva | 17524801 |
| HiLoad® 16/600 Superdex® 75 pg | Cytiva | 28989333 |
| TECAN Spark | TECAN | 30086376 |
| ZipTip® with C18 resin | Merck | ZTC18M096 |
| ESI-LTQ ORBITRAP XL spectrometer | Thermo Scientific | N/A |
| Shigemi 5-mm symmetrical NMR microtube | Shigemi, Co., Ltd. | BMS-005B |
| Bruker INOVA 600 MHz NMR spectrometer | Bruker | N/A |
| Software and Algorithms |  |  |
| TopSpin 4.2.0. | https://www.bruker.com/en/products-and-solutions/mr/nmr-software/topspin.html |  |
| POKY<sup>9</sup> | https://sites.google.com/view/pokynmr |  |
| NMRFAM SPARKY<sup>10</sup> | https://nmrfam.wisc.edu/nmrfam-sparky-distribution/ |  |
| Magellan™ v 7.2 | https://lifesciences.tecan.com/software-magellan |  |
| GraphPad Prism | https://www.graphpad.com |  |
| Microsoft Excel | https://www.microsoft.com/en-us/microsoft-365/excel |  |

### Materials and equipment

#### Loading / Washing Buffer at pH 7.5 (for immobilized metal affinity chromatography, IMAC)

Materials and equipment Loading / Washing Buffer at pH 7.5 (for immobilized metal affinity chromatography, IMAC)
| Reagent | Final concentration | Amount for 1 L |
| --- | --- | --- |
| HEPES | 50 mM | 11.92 g |
| NaCl | 300 mM | 17.53 g |
| Imidazole | 10 mM | 0.68 g |
| Dithiothreitol (DTT) | 1 mM | 0.1543 g |
| ZnCl<sub>2</sub> 1 M | 1 µM | 1 µL |

Prepare 1 L, adjust to pH 7.5, and store at 4 °C for up to one week. Pass through 0.22 μm filter and completely degas before use.

#### Elution Buffer at pH 7.5 (for IMAC)

| Reagent | Final concentration | Amount for 0.5 L |
| --- | --- | --- |
| HEPES | 50 mM | 5.96 g |
| NaCl | 300 mM | 8.76 g |
| Imidazole | 250 mM | 8.52 g |
| DTT | 1 mM | 0.0771 g |

Prepare 0.5 L, adjust to pH 7.5, and store at 4 °C for up to one week. Pass through 0.22 μm filter and completely degas before use.

#### Size Exclusion Chromatography (SEC) Buffer at pH 7.5

| Reagent | Final concentration | Amount for 1 L |
| --- | --- | --- |
| HEPES | 50 mM | 11.92 g |
| NaCl | 100 mM | 5.85 g |
| DTT | 5 mM | 0.7713 g |

Prepare 1 L, adjust to pH 7.5, and store at 4 °C for up to one week. Pass through 0.22 μm filter and completely degas before use.

#### Fluorescence Assay Buffer at pH 7.5

| Reagent | Final concentration | Amount for 0.1 L |
| --- | --- | --- |
| HEPES | 50 mM | 1.19 g |
| DTT | 1 mM | 0.0154 g |
| Triton X-100 | 0.01% (w/v) | 10 µL |

Prepare 0.1 L, adjust to pH 7.5, and store at 4 °C

## Step-by-step method details

### Expression and purification of SARS-CoV-2 PL^pro^

**Timing: 4 days**

In this section, we describe in detail the steps needed to express His_6_-SUMO-PL^pro^ from BL21(DE3) *Escherichia coli* cells harboring the pGTM_CoV2_NSP3_003_SUMO (or pGTM_SUMO_PLpro_C111S) plasmid and subsequently obtain pure PL^pro^ (or C111S-PL^pro^).

1. Expression of His_6_-SUMO-PL^pro^ by induction with IPTG of transformed BL21(DE3) cells (days 1 and 2):

**Note**: Cell culture transfers, reagent additions, and cell sample collection for measuring cellular growth must be carried out under aseptic conditions inside the laminar flow cabinet or using a Bunsen burner to prevent cell culture contamination.

a. Prepare a starter culture using transformed BL21(DE3) cells by selecting a single colony (from plates prepared in step 7 of ***Preparation of LB medium for bacterial growth*** section) in 50 mL of LB broth with 100 μg mL^-1^ ampicillin added. Incubate this starter culture for 16 hours at 37 °C under agitation (at 200 rpm).
b. Transfer 25 mL of the starter culture to 1 L of LB broth with 100 μg mL^-1^ ampicillin added and incubate the suspension at 37 °C under agitation (200 rpm).
c. Measure the optical density at 600 nm (OD_600_) every 30 minutes until it reaches *ca*. 0.6 (approximately 1.5 – 2 hours).
d. Add IPTG and ZnCl_2_ to a final concentration of 1 mM and 1 μM, respectively, to induce protein expression.
e. Incubate for 16-18 hours at 17 °C under constant agitation (200 rpm).

**Alternative:** To prepare isotopically-enriched samples for NMR experiments using MJ9 minimal medium optimized for isotope-enrichment,^6^ at the end of point a above, transfer 25 mL of the obtained starter culture to 1 L of MJ9 minimal medium added with 50 μg mL^-1^ ampicillin. The resulting suspension should be incubated at 37 °C with shaking (200 rpm) and treated as described from point 1c onwards.

**Alternative:** To prepare ^19^F-Trp93 and ^19^F-Trp106 -labeled samples for NMR experiments, use the method for isotopically enriched samples and add 1 aliquot of 5-Fluoroindole concurrently with the IPTG and ZnCl_2_ in step 1d.

2. Cell lysis and soluble extract recovery (Day 3):
  a. Centrifuge the culture for 50 minutes at 3,488 RCF at 4 °C to harvest the cells.
  b. To 30 mL of lysis buffer, add 1 μL of nuclease mix and gently resuspend the harvested cells.
  c. Lyse cells by sonication with microtip, Sonicator S-4000 from Misonix Ultrasonic Liquid Processors. For samples > 30 mL, use amplitude 20 %, process time 15 minutes, time on = 5 sec, time off = 5 sec; for samples < 30 mL, use amplitude = 40 %, process time 15 mins, time on = 5 sec, time off = 5 sec. Keep the pellet on ice for the sonication’s duration and ensure that the tip stays in the center of the sample without touching the vial.
  d. Collect a 15 μL aliquot of the lysate (total lysate, TL) in a microcentrifuge tube for SDS-PAGE analysis; add the appropriate amount of loading dye (usually 5 μL, or what the manufacturer recommends) and freeze at -20 °C.
  e. To separate the soluble extract from the insoluble proteins and cell debris, centrifuge the lysate for 50 minutes at 27,000 × *g* at 4 °C; collect the soluble extract. Collect 15 μL aliquots of each soluble extract (S) and the insoluble pellet (I) for SDS-PAGE analysis, add loading dye to each tube and freeze at -20 °C. Discard the remaining insoluble portion according to standard biosafety protocols.
  f. Filter the soluble extract using 0.8 μm syringe filters and store at 4 °C for purification.

**Critical**: The cell lysate must be kept on ice throughout the procedure to avoid thermal protein denaturation caused by an abrupt temperature increase during sonication.

3. Immobilized metal affinity chromatography (IMAC) (Day 3):

**Note:** The expressed SARS-CoV-2-PL^pro^ contains a His_6_-SUMO tag at its N-terminus, allowing for protein purification using immobilized Ni^2+^ affinity chromatography carried out using an ÄKTApure system with a 5 mL HisTrap HP column (Cytiva) working at a flow rate of 4 mL min^-1^.

**Critical:** SARS-CoV-2-PLpro’s enzymatic activity is partially inhibited at low temperatures. All purification steps described in this protocol were performed on an ÄKTApure system in a cold box at 4 °C.

a. Equilibrate the column with 5 CVs of washing buffer.
b. Load the soluble extract onto the column using a sample pump and collect the unbound material (hereafter called Flow-Through, FT).
c. Wash the column with 20 CVs of washing buffer or until the absorbance at 280 nm (Abs_280_) returns to the baseline to ensure complete removal of the unbound material
d. Collect a 15 μL aliquot of FT for SDS-PAGE analysis, add the loading dye and freeze at - 20 °C. Store the FT at 4 °C.
e. Elute the bound proteins by increasing the imidazole concentration by setting a two-step elution, with the first step at 10 % elution buffer (90 % washing buffer) for 10 CVs and the second step at 100 % elution buffer for 10 CVs. Collect the eluate in 2 mL fractions. A representative chromatogram is shown in **Figure 2A**.
f. Collect a 15 μL aliquot of each fraction that contains protein for SDS-PAGE analysis, add the loading dye and freeze at -20 °C.
g. Analyze the total lysate (TL), the soluble extract (S), as well as the flow-through (FT) and the elution samples from the IMAC for protein presence and purity by running an SDS-PAGE loading 5 μL of the samples previously collected and boiled for 5 minutes at 90 °C (a representative protein gel is shown in **Figure 2B**).
h. Pool the fractions containing His_6_-SUMO-PL^pro^ and store them at 4 °C.

4. Cleavage of the His_6_-SUMO tag by His_8_-MBP-Ulp1-SUMO-protease and buffer exchange (Day 3):
  a. Add 1 aliquot of His_8_-MBP-Ulp1-SUMO protease to the pool of fractions obtained at the end of step 3.
  b. Dialyze the solution containing the His_6_-SUMO-PL^pro^ and the SUMO^pro^ in washing buffer for 16-18 hs at 4 °C.
  c. Collect a 15 μL aliquot for SDS-PAGE analysis, add the loading dye and freeze at -20 °C.
  d. Collect also a 15 μL aliquot of the dialysate for SDS-PAGE analysis (**Figure 2B)**.
  e. Store at 4 °C for purification.

**Figure 2:**
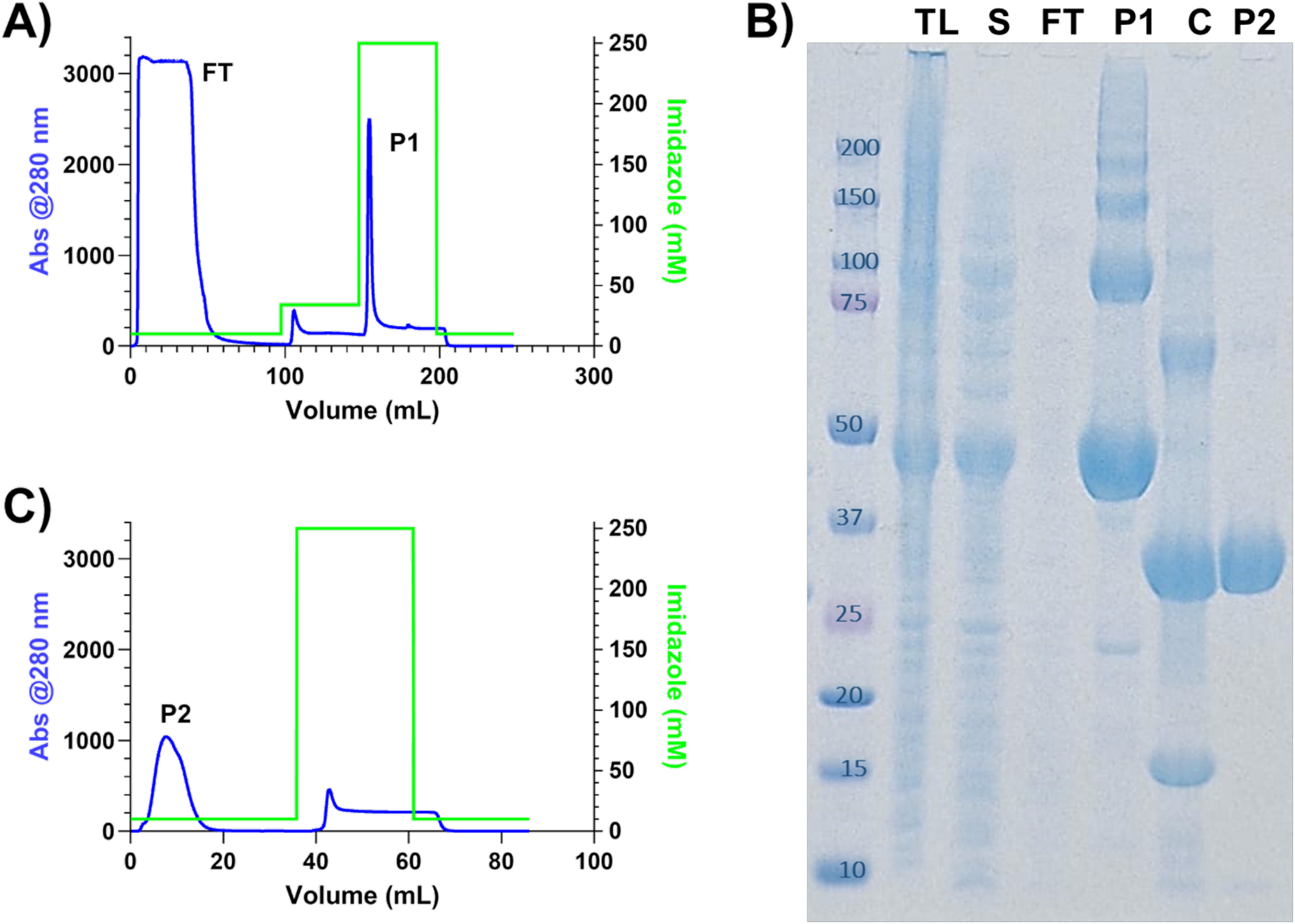
Immobilized metal (nickel) affinity chromatography (IMAC). **(A)** Representative chromatogram showing the entire process of protein loading, flowthrough collection, and protein elution. **(C)** SDS – PAGE showing total lysate (TL), soluble extract (S), flowthrough (FT), fractions eluted in peak 1 (P1) the product after the SUMO-tag cleavage (C), and fractions eluted in peak 2 (P2). His_6_-SUMO-PL^pro^ (molar mass of *ca*. 50 kDa) is detected in lanes corresponding to peak total lysate, soluble extract and P1, a portion of the construct was already cleaved by native proteases, resulting in signals around 37 kDa (PL^pro^) and around 15 kDa (SUMO-tag). In the C lane is possible to detect the SUMO^pro^ (∼70 kDa) and the SUMO-tag (∼15 kDa).

**Note:** The proteolytic cleavage of the N-terminal His_6_-SUMO tag is performed by adding His_8_-MBP-Ulp1-SUMO^pro^ to the expressed His_6_-SUMO-PL^pro^. General protocols for SUMO protease cleavage recommend carrying out cleavage in the presence of imidazole concentrations lower than 150 mM to prevent the adverse effects of imidazole on the activity of SUMO^pro^. Therefore, the cleavage performed in dialysis is useful in decreasing the concentration of imidazole and can be performed with Washing Buffer without imidazole.

**Note**: Incubation times for complete cleavage of the N-terminal His_6_-SUMO tag by His_8_-MBP-Ulp1-SUMO^pro^ may vary depending on the protease activity. In this protocol, a 1:100 (w/w) ratio of SUMO^pro^ generally provides > 99 % cleavage after 3 hours when carried out in a pH 7.5 buffer containing a reducing agent (*e*.*g*., the Elution Buffer) at 26 °C.

**Note**: The His_8_-MBP-Ulp1-SUMO protease is aliquoted at 1 mg mL^-1^ and stored at -80 °C in a buffer containing 50 mM Tris-HCl, 150 mM NaCl, 10 % glycerol, and 0.5 mM DTT, at an adjusted pH of 7.5. The protease is used at a ratio (w/w) of 1:1000 to 1:100 to cleave the His_6_-SUMO tag from His_6_-SUMO-PL^pro^ to obtain native PL^pro^.^5^

5. Removal of the cleaved tag and SUMO protease from cleaved PL^pro^ solution (Day 4):

**Note:** The cleaved His_6_-SUMO tag and His_8_-MBP-Ulp1-SUMO-protease present N-terminal His-tags, allowing for separation from the native PL^pro^ through an additional step of IMAC. The cleaved PL^pro^ will not interact with the Ni resin and will be collected in the flowthrough. The His_6_-SUMO tag and His_8_-MBP-Upl1-SUMO protease will bind to the column and be eluted at a higher imidazole concentration. Separation is carried out using an ÄKTA Pure system and a 5 mL HisTrap HP column (Cytiva) loaded with Ni^2+^, working at a flow rate of 4 mL min^-1^.

a. Wash a 5 mL HisTrap HP column (Cytiva) with 5 CVs of MilliQ water and equilibrate it with 5 CVs of Loading / Washing Buffer.
b. Load the protein solution onto the column while using the Loading / Washing Buffer while collecting the flow through in the fraction collector.
c. Wash the column with Loading / Washing Buffer while monitoring the absorbance at 280 nm (Abs_280_) to fractionate the cleaved, native PL^pro^. Store the fractions containing the protein solution at 4 °C.
d. Collect a 15 μL aliquot of protein solution in a microcentrifuge tube for SDS-PAGE analysis, add the loading dye, and freeze at -20 °C (**Figure 2C**).
e. Elute the bound proteins remaining on the column using 5 CVs of 100 % Elution Buffer until the Abs_280_ baseline returns to zero, which will happen around 5 CVs.
f. Collect a 15 μL aliquot of the eluted proteins in a microcentrifuge tube for SDS-PAGE analysis, add the loading dye and freeze at -20 °C (**Figure 2C**).

6. Size exclusion chromatography (SEC) (Day 4):

**Note:** A final size exclusion chromatography (SEC) purification step provides a pure and monodisperse PL^pro^ sample in a buffer suitable for downstream structural and biochemical studies. This step is carried out using an ÄKTA Pure system set up at 4 °C and a HiLoad® 16/600 Superdex® 75 pg column (Cytiva), working at a flow rate of 0.8 mL min^-1^.

a. Concentrate the protein solution to a final volume 2-5 mL using a Sartorius Vivaspin PES centrifugal concentrator (MWCO 10,000 Da) (Sartorius).

**Critical:** Although not strictly necessary, this concentration step is highly recommended to expedite the following SEC. During this step, it is important to monitor the concentration of the protein solution (by measuring the absorbance at 280 nm and considering ε_280_ = 45,270 M^-1^ cm^-1^), which should not exceed 0.2 mM to avoid possible precipitation. If the desired volume of 2-5 mL is not achievable, split the protein solution into two or more aliquots to be independently treated in the next purification step.

b Wash the column with one CV of MilliQ water and equilibrate it with at least 1.35 CVs of SEC buffer.
c Load up to 5 mL of protein solution using a sample loop of the appropriate volume and carry out the protein elution using SEC buffer (a representative chromatogram is shown in **Figure 3A**), collecting the eluate in 2 mL fractions and store at 4°C.
d Collect a 15 μL aliquot of each fraction in microcentrifuge tubes for SDS-PAGE analysis, add the loading dye, and freeze at -20 °C.
e Repeat points c - d for the remaining protein solution.
f Analyze all collected fractions for protein presence and purity by running an SDS-PAGE gel loading 7 μL of the samples previously collected and boiled for 5 minutes at 90 °C. Pool the fractions containing native PL^pro^. A representative SDS-PAGE is shown in **Figure 3B**.
g Native PL^pro^ can be concentrated up to 6.5 mg mL^-1^ (corresponding to 0.14 mM). At these concentrations, native PL^pro^ can be flash-frozen in liquid nitrogen and stored for up to one month at -80 °C.

**Figure 3:**
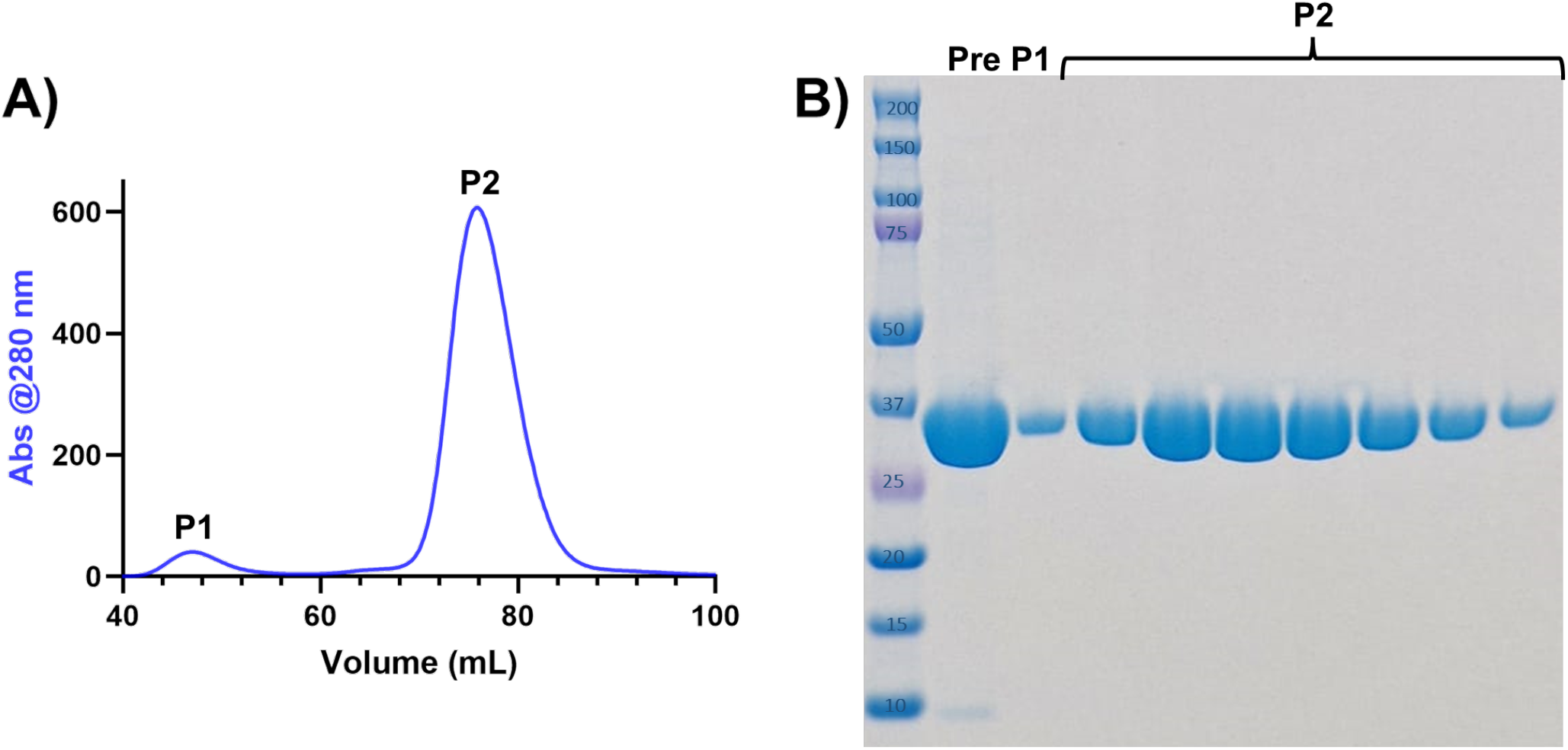
Size exclusion chromatography of native PL^pro^. **(A)** Representative SEC chromatogram and **(B)** corresponding SDS – PAGE from elution of the SEC column. Lane “Pre” corresponds to the pre-loaded cleaved protein solution, followed by native PL^pro^ eluted in peak 1, which is a small portion of dimer, and peak 2, for the pure protein (at a molar mass of *ca*. 37 kDa).

**Note:** PL^pro^ should be frozen in aliquots of appropriate concentration and volume for their respective experiments for isotopically enriched samples. Generally, enzymes should be flash frozen in 50-100 μL aliquots to minimize the need for thaw/freeze cycles and to ensure rapid freezing. For the ^19^F-Trp labeled samples, the protein can be concentrated to 50 μM and stored in 270 μL aliquots. For ^15^N-labeled samples, the protein can be concentrated to 100 μM and stored in 270 μL aliquots.

**Note:** The SEC buffer reported here is one of several buffers where native PL^pro^ can be stored. Buffers most compatible with other downstream studies can also be chosen.

### Characterization of SARS-CoV-2 PL^pro^ by mass spectrometry

**Timing: 3 h**

7. Sample preparation by ZipTip®:

**Note:** 10 **-** 20 μL of pure protein, at least 50 μM concentration, is used for the chromatography, following the manufacturer’s directions (here reported).

**Critical:** Ensure samples are within the instrument’s detection limits. Generally, a good MS signal should be obtained with 1 picomole of the sample.

a. Adjust sample to 0.1 % trifluoroacetic acid (TFA); final sample pH should be < 4.
b. Depress the pipettor plunger to a dead stop. Using the maximum volume setting of 10 μL, aspirate the wetting solution (100 % acetonitrile) into the tip.
c. Dispense to waste. Repeat step b.
d. Aspirate equilibration solution (0.1 % TFA in Milli-Q® solution grade water).
e. Dispense to waste. Repeat step d.
f. Bind the sample to the ZipTip® pipette tip by fully depressing the plunger to a dead stop.
g. Aspirate and dispense the sample 7–10 cycles for maximum binding of complex mixtures.
h. Aspirate wash solution (0.1 % TFA in Milli-Q grade water) is dispensed to the tip and dispensed to waste.
i. Repeat step h at least once. **Note**: A 5 % methanol in 0.1 % TFA/water wash can improve desalting efficiency.
j. With a normal tip, dispense 1 to 4 μL of elution solution (1 % formic acid / 50 % methanol) into a clean vial using a standard pipette tip. **Critical**: Acetonitrile and methanol are volatile, and evaporation can occur rapidly. If this occurs, add more eluant to recover the sample.
k. Carefully aspirate and dispense eluant through the ZipTip pipette tip at least three times without introducing air into the sample. **Note:** Sample recovery can be improved (at the expense of concentration) by increasing the elution volume to 5 μL.

8. Analyze the eluted species using an ESI-MS spectrometer (Thermo Scientific LTQ ORBITRAP XL) using the software provided with the instrument.

Note: The resulting deconvoluted spectrum of pure protein, **Figure 4A and B**, shows a single signal corresponding to an experimental molar mass of 35,934 Da. The experimental value perfectly matches that estimated by the amino acid protein sequence and establishes that the current protocol for the expression and purification of SARS-CoV-2 PL^pro^ is a reliable procedure to obtain large quantities of native PL^pro^.

**Figure 4:**
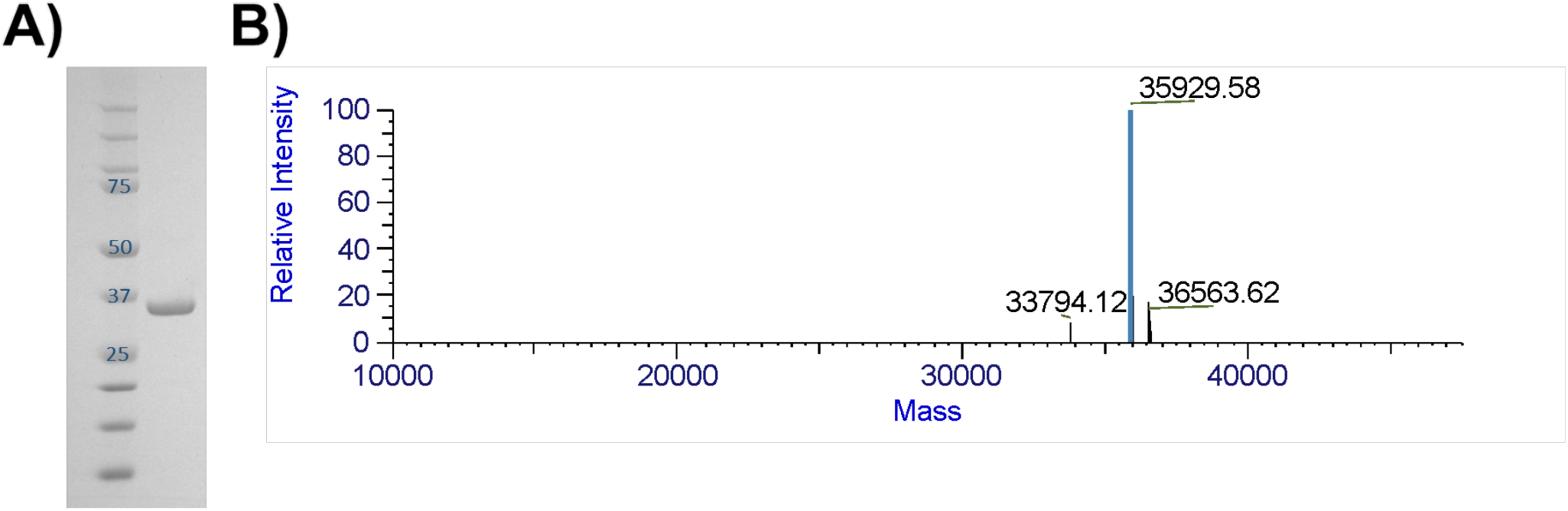
Characterization of purified PL^pro^. **(A)** SDS-PAGE gel showing the high quality of SARS-CoV-2 PL^pro^ sample. Lane a – molecular weight standards; Lane b – SARS-CoV-2 PL^pro^ post-SEC purification (sample prepared as described above) **(B)** ESI-MS profile of SARS-CoV-2 PL^pro^ showing the signal corresponding to an experimental molecular mass of 35929.58 Da (Expected mass= 35,928.9 Da).

### ^19^F NMR assay for screening potential PL^pro^ inhibitors

**Timing: 2 h**

**Note**: PL^pro^ contains 2 tryptophan residues (W93 and W106) that can be labeled with ^19^F through the incorporation of 5-Fluoroindole added at the induction step (as detailed in Expression and Purification of SARS-CoV-2 PLpro). The two residues are located in close proximity to the active site, and each provides a unique peak in a 1-D fluorine spectrum, as shown below.

We have demonstrated that known PL^pro^ inhibitors cause both peaks to shift while other small molecules cause no change to the ^19^F spectrum (**Figure 5A-D**). Peaks were assigned by collecting a spectrum on a ^19^F labeled W106F mutant shown as an overlay in **Figure 5B. Figure 5C** shows the effect of GRL0617 compared to the negative control Remdesivir (**Figure 5D**). This assay requires very low concentrations of ^19^F labeled PL^pro^ and potential inhibitors. It, therefore, is a simple screen that can be used to identify potential inhibitors that can be further characterized using the FRET assay described later in this protocol.

**Figure 5:**
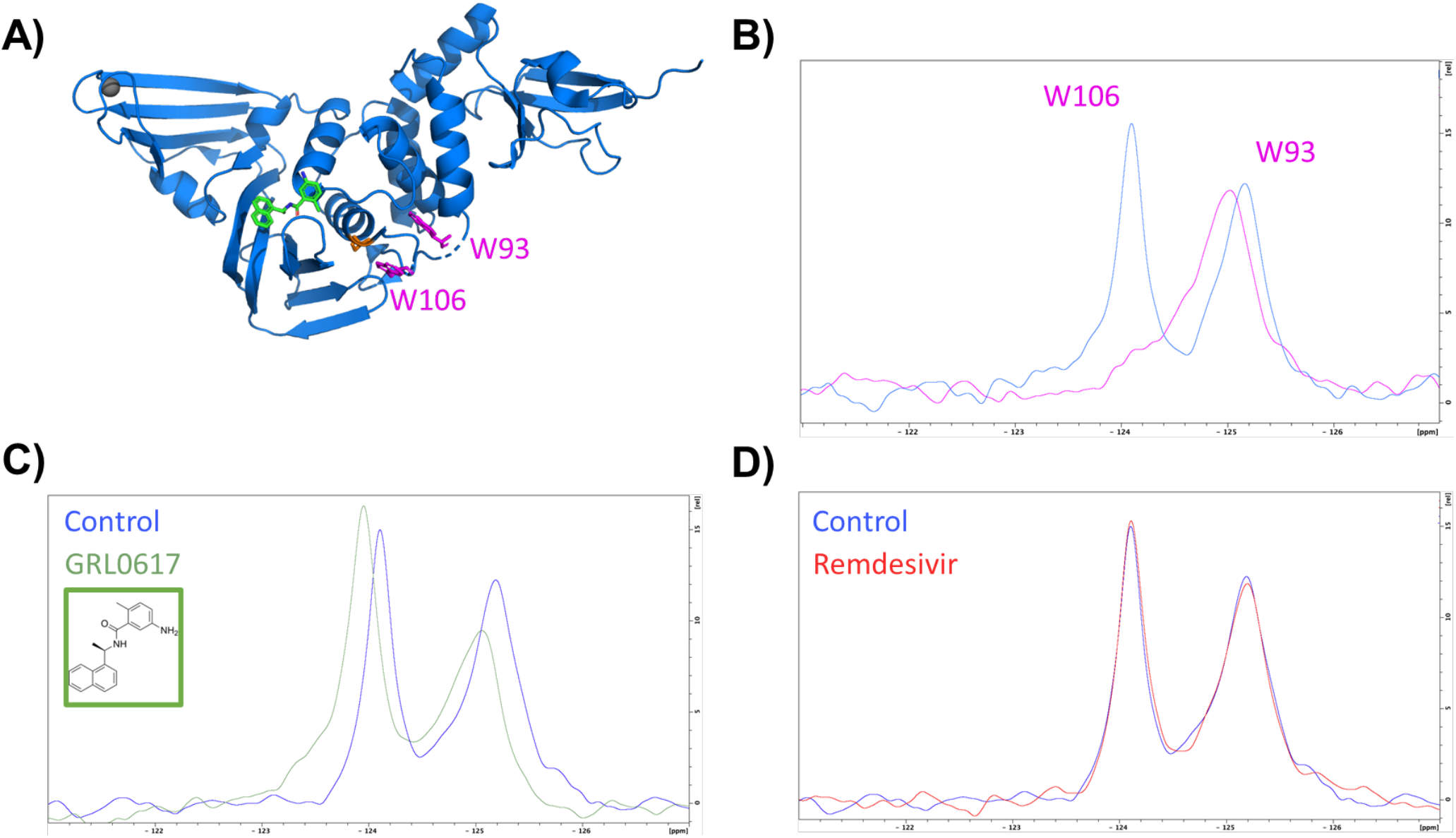
^**19**^**F 1D spectra of the protein. (A)** Pymol structure of PL^pro^ (PDB_ID 7JCM, in blue) bound to the GRL0617 (green)^17^, together with the locations of the active-site Cys111 (red), and two tryptophans at positions 93 and 106 (magenta). **(B)** 1D ^19^F-NMR spectrum of the [^19^F-W93, ^19^F-W106] PL^pro^ shows signals, one for each of the two fluorinated tryptophans. 1D ^19^F-NMR spectra of the PL^pro^ exposed to **(C)** GRL0617 (green spectrum) and **(D)** Remdesevir (red spectrum), compared to the protein alone (control, blue spectra).

9. Sample preparation for the ^19^F NMR experiment:
  a. Prepare 6 mM stock solutions in DMSO of all inhibitors to be used in the screen.
  b. Gently thaw n + 1 of the 270 μL aliquots of purified 0.05 mM ^19^F-Trp labeled PL^pro^ on ice (n = number of inhibitors to be screened).
  c. Add 3 μL of inhibitor to each protein sample.
  d. Add 3 μL of DMSO to the control.
  e. Add 30 μL deuterium oxide (Cambridge Isotopes) to each protein sample, needed for spectrometer lock.
  f. Transfer the samples to Shigemi 5-mm symmetrical NMR microtubes (Shigemi, Co.).
10. Data recording and analysis:
  g Record ^19^F 1D spectra of the protein at 298 K on a Bruker 600 MHz NMR spectrometer using 0.5 increments, 1000 scans, and 0.7 s relaxation delay. A sweep width of 20 ppm and an offset of -125 ppm are used for the ^19^F dimension. The total acquisition time for each experiment is 13.5 minutes.
  h Process and analyze the obtained spectra using Bruker TopSpin (v4.0) (**Figure 5**).

### Fluorescence assay for the characterization of PL^pro^ inhibition by IC_50_ determination

**Timing: 2 h**

**Note:** The inhibitory properties of PL^pro^ are assessed using fluorescence assays with the peptide substrate Z-Arg-Leu-Arg-Gly-Gly-AMC Acetate, which includes the PL^pro^ cleavage site RLRGG/X and is labeled with an AMC fluorophore (CAS #167698-69-3; VWR #I-1690.0100BA or Bachem #4027158.0100). When PL^pro^ cleaves the labeled peptide, an increase in fluorescence is observed at the emission maximum of the AMC (450 nm). This increase in fluorescence is directly proportional to the enzyme activity.

Fluorescence assays are conducted at varying concentrations of potential PL^pro^ inhibitors to screen candidate molecules and determine their IC_50_ values (the concentration of inhibitor that results in a 50% reduction in enzyme activity).

These assays are performed using black opaque 96-well microplates (Corning) with a final volume of 100 μL in each well. Experiments should be conducted in triplicate, and results monitored and analyzed using a TECAN Infinite M1000 microplate reader along with Magellan™ software (or a comparable multi-well fluorimeter).

11. Instrument protocol setup: Use the Magellan™ software of TECAN Infinite M1000 microplate reader with the following parameters: (i) Excitation wavelength: 350 nm; (ii) Emission wavelength: 450 nm. Fluorescence intensity is recorded every 3 seconds for a total of 15 – 60 minutes at 20 °C.

**Note**: Raw data collected during the experiments indicate fluorescence intensity due to substrate proteolysis and are reported in terms of Relative Fluorescence Units (RFU).

**Note:** Depending on the range of concentration of the tested molecule and the inhibition of the enzyme, the fluorimeter must be calibrated to avoid exceeding the instrument’s sensitivity.

12. PL^pro^, substrate, and inhibitor preparation:
  a. Prepare a 500 μM fluorescent substrate stock solution in assay buffer and store it as 60 μL aliquots in PCR-strips tubes at -20 °C.
  b. Gently thaw a small aliquot of purified PL^pro^ on ice.
  c. Estimate the protein concentration using the Lambert-Beer equation, with Ill_280_ = 45,270 A.U. M^-1^ cm^-1^.
  d. Dilute the sample to a final concentration of 208.3 μM using assay buffer, at least 10 mL for each 96-well plate.
  e. Prepare inhibitor stocks, dissolving them in DMSO and diluting them to 100 mM concentrations in DMSO.
  f. Prepare the working solution for the inhibition test. Here, we will report the step-by-step preparation to test seven different concentrations in the range 100 μM to 100 nM in triplicate, in a final DMSO concentration of 1 % (**Figure 6A-E**):
    i. In PCR strip tubes, add 9 μL of pure DMSO in the odd positions (1, 3, 5, 7) and 5 μL in the other tubes (position 2, 4, 6, and 8).
    ii. Take 1 μL of the 100 mM stock solution, add it into the tube at position one, and mix, obtaining 10 μL of a solution at 10 mM.
    iii. Take 1 μL of the 10 mM solution from the previous step, add it into the tube at position three, and mix, obtaining 10 μL of a solution at 1.0 mM (**Figure 6A**).
    iv. Take 1 μL of the 1.0 mM solution from the previous step, add it into the tube at position five, and mix, obtaining 10 μL of a solution at 0.1 mM (100 μM) (**Figure 6A**).
    v. Take 1 μL of the 0.1 mM solution from the previous step, add it into the tube at position five, and mix, obtaining 10 μL of a solution at 0.01 mM (10 μM) (**Figure 6A**).
    vi. Take 5 μL of the 10 mM solution from the first tube, add it into the tube at position two, and mix, obtaining 10 μL of a solution at 5.0 mM (**Figure 6A**).
    vii. Take 5 μL of the 1 mM solution from the third tube, add it into the tube at position four, and mix, obtaining 10 μL of a solution at 0.5 mM (**Figure 6A**).
    viii. Take 5 μL of the 0.1 mM solution from the fifth tube, add it into the tube at position six, and mix, obtaining 10 μL of a solution at 0.05 mM (**Figure 6A**).

**Figure 6:**
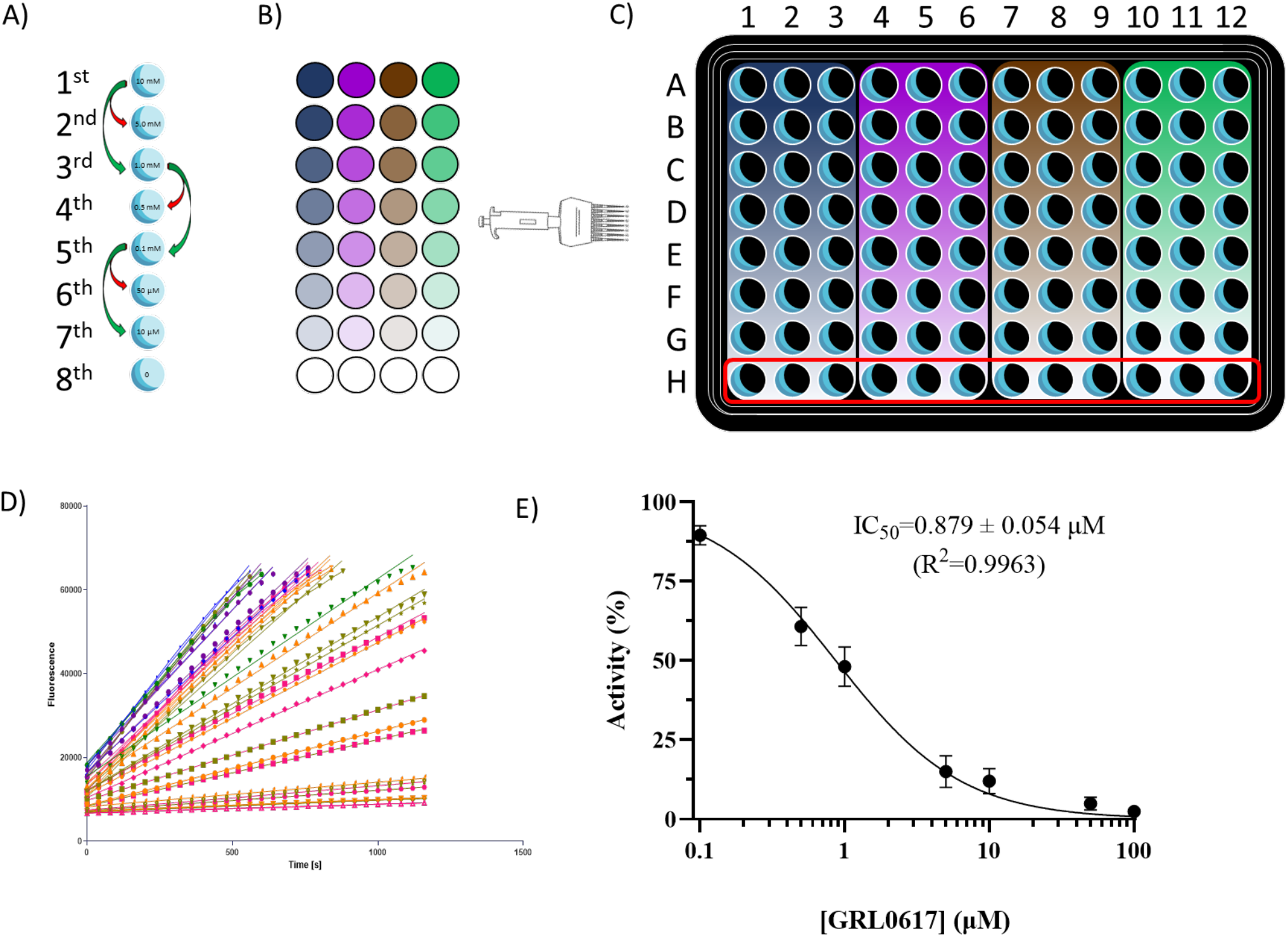
Drugs preparation and data processing for IC_50_ determination. **(A)** Schematic of drug preparations, using the method of serial dilution to cover the range 100 μM - 100 nM in a PCR 8-tube strip. **(B)** Result of the drug preparation for 4 different drugs, to be tested in triplicate using one 96-well plate. **(C)** Representation of the final result of the addition of the drugs to the protein in the plate, stronger colors indicate higher concentrations, whereas faded colors indicate lower concentrations. (D) Representative experimental raw data showing the increase of fluorescence intensity as a function of reaction time. (E) Representative dose-response curve for GRL0617, with error bars representing the S.D.s of triplicate measurements.

**Note:** When preparing the inhibitor(s) stock solution, if possible, it is preferable to dissolve it in deuterated DMSO; this way, the stock solution can be used to perform ^1^H-NMR for qualitative and/or quantitative purposes.

**Note:** Inhibitors vary in their solubility, so the assay may be adjusted to use final concentrations of DMSO as low as 1 % for appropriately soluble substrates and inhibitors.

**Note:** The same tip could be used for steps i to v when preparing the working solution for the inhibition test to decrease the volume lost during preparation.

**Note**: The range for the test can be modified as needed, as it depends on the first tube concentration (**Figure 6A and B**).

13. Setup of the reaction mixtures in the 96-well microplates:
  i. In each well of the 96-well microplate, add 95 μL PL^pro^ solution obtained in step 13d using a multichannel pipette (**Figure 6C**).
  ii. Using an 8-channel micropipette, add 1 μL of the inhibitor working solution for each well in a column, and repeat this for at least 3 columns (**Figure 6B and C**).
  iii. Place the 96-well microplate in the Infinite M1000 plate reader (TECAN) and let the plate equilibrate inside the plate reader for at least 10 minutes at 20 °C, reading the fluorescence during the interaction using the parameter described in step 11
  iv. Using an 8-channel micropipette, initiate the reaction by adding 4 μL of the 500 μM fluorescent substrate (resulting in 20 μM substrate final concentration) and mix by pipetting up and down 2-3 times.
  v. Start the program and read immediately for at least 15 minutes to measure initial enzyme velocities.

**Note:** The enzymatic reaction begins as soon as the substrate is added to each well. Carefully pipette the substrate into each well, moving as quickly as possible to initiate the data collection with minimal dead time.

14. Data export and analysis:
  a. Export the data from Magellan in Microsoft Excel .xls or in .csv format.
  b. Analyze the data using GraphPad or other kinetic analysis software:
    i. Plot the fluorescence intensity *vs*. time data (**Figure 6D**) for each inhibitor concentration.
    ii. Measure the slope of the linear portion of each curve (usually within the first 5 minutes of reaction) by performing a linear fit on that region. Each obtained slope value corresponds to the initial velocity in the presence of that inhibitor concentration (*V*_*i*_). The slope determined for the control measurements in blue H (**Figure 6D**), carried out in the absence of an inhibitor, corresponds to the initial velocity of the non-inhibited PL^pro^ (*V*_*0*_).
    iii. Calculate the average ± standard deviation values for the triplicate measurements carried out at each inhibitor concentration to obtain, for each concentration, an averaged *V*_*i*_ (or *V*_*0*_) value.
    iv. Using the averaged values, calculate residual activity (%) at each inhibitor concentration using the formula:

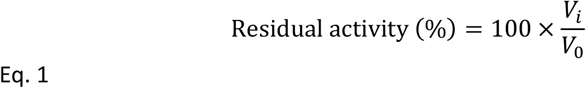
    v. Plot residual activity (%) data as a function of increasing inhibitor concentration and fit them by using the following equation:

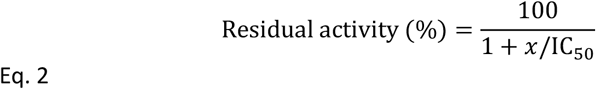
    vi. Transform the *x*-axis visualization on a Log_10_ scale to show the plotted experimental data and resulting fit following the typical sigmoidal behavior (**Figure 6E**). The IC_50_ value estimated by Eq. 2 corresponds to the inhibitor concentration at which residual activity is 50 %.

**Note:** A single 96-well plate can accommodate triplicates for 4 different compounds, and line H will contain only controls (wells with enzyme and substrate in buffer containing 1 % of DMSO). To have a more robust *V*_*0*_, the average of the full line (12 wells) should be used.

**Note**: In case of compounds with intrinsic fluorescence at the assay setting, discount the average fluorescence recorded during the interaction to the specific curves, *i*.*e*. discount the average of the fluorescence for the specific compound concentration to the respective curve obtained after adding the substrate to the solution.

### NMR spectroscopy of PL^pro^

**Timing: Approximately 4 h**

15. Sample preparation for the ^15^N NMR experiment:
  a. Gently thaw a 270 μL aliquot of purified 0.2 mM ^15^N-enriched PL^pro^ on ice.
  b. Add 30 μL deuterium oxide (CortecNet) to the protein solution and transfer the clear, transparent sample to the Shigemi 5-mm symmetrical NMR microtube (Shigemi, Co.).

16. Data recording and analysis:
  a. Record ^1^H,^15^N BEST-TROSY spectra^13-15^ of the protein at 298 K on a Bruker 600 MHz NMR spectrometer using 256 increments, 4 scans, and 1.0 s relaxation delay. Sweep widths of 18 ppm and 40 ppm and offsets of 4.7 ppm and 117 ppm are used for the ^1^H and ^15^N dimensions, respectively. The total acquisition time for each experiment is 135 minutes
  b. Process the spectra using the Bruker TopSpin (v4.0).
  c. Analyze the obtained spectra (**Figure 7**) using the POKY^9^ or NMRFAM SPARKY^10^.

**Figure 7:**
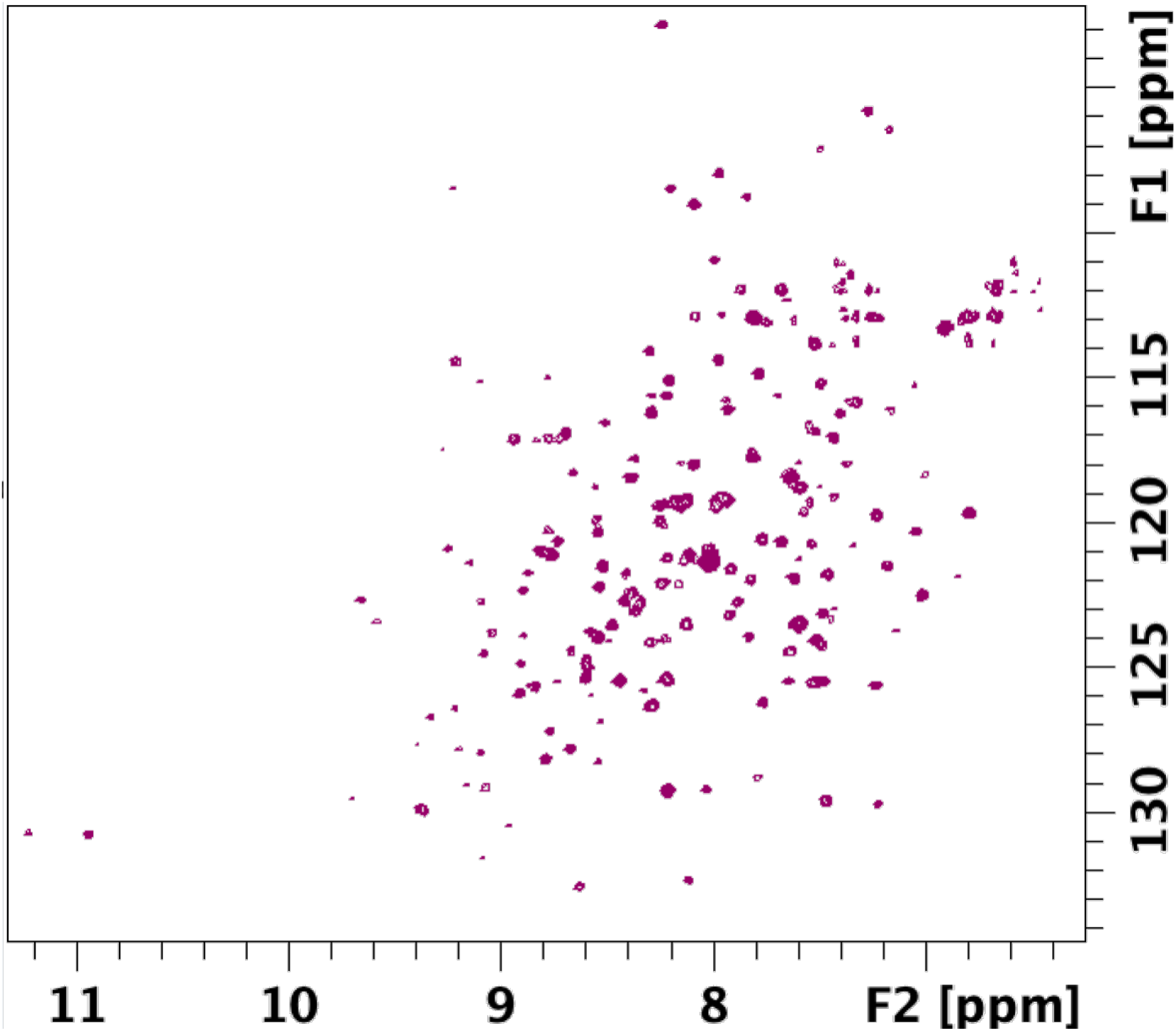
2D ^1^H,^15^N BEST-TROSY NMR spectrum of SARS-CoV-2 PL^pro^ at 298 K. No significant sample degradation was observed over 3 days at room temperature.

**Alternative:** Alternative software for processing NMR data includes NMRPipe, earlier versions, the CcpNmr software suite, and software available in the NMRBox collection^11,12^.

Partial sequence-specific resonance assignments for SARS-CoV-2 PL^pro^ are available in the BMRB entry 51992^16^ and can be used to validate the spectrum and to label the observed peaks following the nearest neighbor criterium.

### Quality control by specific activity measurements of SARS-CoV-2 PL^pro^

If available, sample quality control includes crystallography and NMR data, SDS-PAGE, high-resolution ESI-MS, and specific enzyme activity measurements. The specific activity of the PL^pro^ enzyme was determined to be 11.8 ± 0.557 nmol of substrate cleaved per minute per milligram of enzyme, assayed using 0.2 μM PL^pro^ and 20 μM Z-Arg-Leu-Arg-Gly-Gly-AMC Acetate in fluorescence assay buffer (described in the Preparation of reagent stock solutions), for a final volume per well of 100 μL (considering the extinction coefficient of Z-Arg-Leu-Arg-Gly-Gly-AMC Acetate at 460 nm in 50 mM HEPES and 1 mM DTT is approximately 1,800 M^−1^cm^−1^).

### Expected outcomes

This paper provides a detailed protocol for the expression, purification, and production of tens of milligram quantities of highly pure (> 99 %) SARS-CoV-2 PL^pro^ with native N and C termini and in a biochemically active form, as demonstrated by the biostructural characterization and activity assays reported. The yields of pure PL^pro^ obtained depend on the medium used for fermentation.

PL^pro^ samples produced by this protocol are stable in NMR measurements for days at room temperature and provide excellent ^19^F-NMR and [^15^N-^1^H]-HSQC-TROSY spectra suitable for structure, dynamic, and ligand binding studies. The paper also presents a standardized assay for estimating IC_50_ for new potential PL^pro^ inhibitors, thus complementing structural information with biochemical insights into the catalytic and inhibition mechanisms of this target.

**Table 4:** Yields of pure native PL^pro^ obtained in different media. Protein was spectrophotometrically quantified assuming MW = 35,976 Da and ε_280_= 45,270 A.U. M^-1^ cm^-1^).

| Medium | Yield (mg L<sup>-1</sup>) |
| --- | --- |
| LB | 15 - 38 |
| MJ9 (minimal medium) | 5 - 14 |

### Limitations

The protocol section that describes the characterization of enzyme inhibition by fluorescence assays assumes solubility of the screening molecules in 1 % DMSO. For potent inhibitors which can be studied at lower concentrations, lower DMSO concentrations may be used if solubility allows. Each potential inhibitor should be evaluated for solubility prior to setting up the assay. Moreover, IC_50_ obtained from the biochemical assays is a preliminary measurement of inhibitor potency. A more accurate investigation of the inhibition could be provided by determining thermodynamic and kinetic parameters for the inhibition mechanism.

## Troubleshooting

### Problem 1

Following the 12 - 16 hours of cell growth on LB Petri dishes, no colonies are visible.

### Potential solution

After checking that each step of the transformation procedure was carried out correctly (e.g., correct antibiotic, plasmid concentration, duration of heat shock), the concentration of the cell suspension can be increased via centrifugation. At the end of section **Error! Reference source not found.Error! Reference source not found**. in the “***Before you begin***” section, centrifuge the cell suspension for 1 min at 2500 × *g* and decant half of the supernatant. Resuspend the pellet in the remaining supernatant for a more concentrated cell culture.

### Problem 2

During cell lysis (step 2. Cell lysis and soluble extract recovery), heat generated by sonication and can cause aggregation or misfolding of the enzyme, especially in the enzymatically active form.

### Potential solution

Sonication should be done in short blasts, with the sample cooled in an ice bath in a container with good thermal conductivity. A microfluidizer or homogenizer can be used in place of the aforementioned techniques.

### Problem 3

Uncontrolled cleavage of the His_6_-SUMO tag from His_6_-SUMO-PL^pro^, probably due to trace amounts of native *E. coli* proteases, has been observed at the end of step 3 (IMAC), just before the addition of SUMO^pro^.

### Potential solution

Uncontrolled cleavage can be minimized by carrying out cleavage in imidazole immediately after purifying the His_6_-SUMO-PL^pro^, with little or no negative effects on the production of pure and native PL^pro^, or by on-column cleavage. These protocols, and working at 4 °C as outlined above, can minimize uncontrolled proteolytic cleavage during purification.

### Problem 4

Incomplete cleavage of the His_6_-SUMO-PL^pro^ fusion is observed (*i*.*e*., full target fusion is visible in the final elution from the IMAC).

### Potential solution

Collect the elution fractions containing the His_6_-SUMO-PL^pro^ fusion, repeat the dialysis/buffer exchange and simultaneous cleavage in the presence of His_8_-MBP-Ulp1-SUMO^pro^, and repeat an IMAC step by collecting the flow-through.

### Problem 5

After combining enzyme and substrate during the fluorescence assay control (described in step 14), no increasing fluorescence signal is observed.

### Potential solution

PL^pro^ is very sensitive to oxidation and can lose enzymatic activity due to oxidation of the active site cysteine, even though reducing agents are present. It is suitable to work always with fresh and aliquots. During purification, it is advantageous to work quickly and at 4 °C as much as possible to minimize this effect. Samples can also be protected from oxidation by storing them under nitrogen or argon gas.

### Problem 6

The quality of [^15^N-^1^H]-HSQC-TROSY spectrum is not optimal.

### Potential solution

Prepare a ^2^H,^15^N -enriched sample to obtain a higher quality [^15^N-^1^H]-HSQC-TROSY spectrum.

## Resource availability

### Lead contact

Further information and requests for resources and reagents should be directed to and will be fulfilled by the lead contact, G.T. Montelione.

### Materials availability

Plasmids generated in this study have been deposited to AddGene, and are available under the Uniform

Biological Material Transfer Agreement (“UBMTA”):

Plasmid pGTM_COV2_NSP3_003_SUMO. AddGene ID: 233739

Plasmid pGTM_SUMO_PLpro_C111S. AddGene ID: 233740

Plasmid pGTM_ SUMO-N-StrepII-PLpro. AddGene ID: 234322

Plasmid pGTM_ SUMO-N-8XHis-PLpro. AddGene ID: 233803

Plasmid pGTM_ SUMO-C-8XHis-PLpro. AddGene ID: 234321

Plasmid pGTM_ SUMO-N-8XHis-PLpro-C111S. AddGene ID: 234472

Plasmid pGTM_SUMO_PLpro_W93F_001. AddGene ID: 234490

Plasmid pGTM_SUMO_PLpro_W106F_001. AddGene ID: 234491

Plasmid pGTM_YR375_SUMO_Protease_001. AddGene ID: 190063

## Acknowledgments

We thank L. Spaman, E. Gormley and Dr. A. Gaur, for helpful discussions, suggestions, and comments on the manuscript. This research was supported by National Institutes of Health grant 1U19 AI171443 (to S. Chanda, A. García-Sastra, and G.T.M.). R.G.-C. was supported in part by a NIGMS Graduate Training Grant (T32-GM141865). This work was made possible by the RPI Core Facilities (Microbiology & Fermentation, Proteomics & NMR) and NIH Shared Instrumentation Award S10-OD030482.

## Author contributions

These authors contributed equally as first authors: A.D.F.. and R.G.-C. Conceptualization: A.D.F., R.G.-C., and G.T.M. Biochemical investigations: A.D.F., R.G.-C., B.S. and S.Z., NMR investigations: A.D.F., R.G.-C., B.S. and T.A.R. L.M. Visualization: A.D.F., R.G.-C., B.S., S.Z., T.B.A., T.A.R. and G.T.M. Writing, reviewing, and editing: A.D.F., R.G.-C., B.S., S.Z., T.B.A., T.A.R. and G.T.M. Funding acquisition and supervision: G.T.M.

## References

1. Osipiuk, J., Azizi, S.A., Dvorkin, S., Endres, M., Jedrzejczak, R., Jones, K.A., Kang, S., Kathayat, R.S., Kim, Y., Lisnyak, V.G., et al. (2021). Structure of papain-like protease from SARS-CoV-2 and its complexes with non-covalent inhibitors. Nat Commun 12, 743. 10.1038/s41467-021-21060-3.

2. Gao, X., Qin, B., Chen, P., Zhu, K., Hou, P., Wojdyla, J.A., Wang, M., and Cui, S. (2021). Crystal structure of SARS-CoV-2 papain-like protease. Acta Pharm Sin B 11, 237–245. 10.1016/j.apsb.2020.08.014.

3. Chan, H.T.H., Brewitz, L., Lukacik, P., Strain-Damerell, C., Walsh, M.A., Schofield, C.J., and Duarte, F. (2024). Studies on the selectivity of the SARS-CoV-2 papain-like protease reveal the importance of the P2’ proline of the viral polyprotein. RSC Chem Biol 5, 117–130. 10.1039/d3cb00128h.

4. van Huizen, M., Bloeme-Ter Horst, J.R., de Gruyter, H.L.M., Geurink, P.P., van der Heden van Noort, G.J., Knaap, R.C.M., Nelemans, T., Ogando, N.S., Leijs, A.A., Urakova, N., et al. (2024). Deubiquitinating activity of SARS-CoV-2 papain-like protease does not influence virus replication or innate immune responses in vivo. PLoS Pathog 20, e1012100. 10.1371/journal.ppat.1012100.

5. Mazzei, L., Greene-Cramer, R., Bafna, K., Jovanovic, A., De Falco, A., Acton, T.B., Royer, C.A., Ciurli, S., and Montelione, G.T. (2023). Protocol for production and purification of SARS-CoV-2 3CL(pro). STAR Protoc 4, 102326. 10.1016/j.xpro.2023.102326.

6. Jansson, M., Li, Y.C., Jendeberg, L., Anderson, S., Montelione, G.T., and Nilsson, B. (1996). High-level production of uniformly (1)(5)N- and (1)(3)C-enriched fusion proteins in Escherichia coli. J Biomol NMR 7, 131–141. 10.1007/BF00203823.

7. Bertani, G. (2004). Lysogeny at mid-twentieth century: P1, P2, and other experimental systems. J Bacteriol 186, 595–600. 10.1128/JB.186.3.595-600.2004.

8. Acton, T.B., Xiao, R., Anderson, S., Aramini, J., Buchwald, W.A., Ciccosanti, C., Conover, K., Everett, J., Hamilton, K., Huang, Y.J., et al. (2011). Preparation of protein samples for NMR structure, function, and small-molecule screening studies. Meth Enzymol 493, 21–60. 10.1016/B978-0-12-381274-2.00002-9.

9. Lee, W., Rahimi, M., Lee, Y., and Chiu, A. (2021). POKY: a software suite for multidimensional NMR and 3D structure calculation of biomolecules. Bioinformatics 37, 3041–3042. 10.1093/bioinformatics/btab180.

10. Lee, W., Tonelli, M., and Markley, J.L. (2015). NMRFAM-SPARKY: enhanced software for biomolecular NMR spectroscopy. Bioinformatics 31, 1325–1327. 10.1093/bioinformatics/btu830.

11. Skinner, S.P., Fogh, R.H., Boucher, W., Ragan, T.J., Mureddu, L.G., and Vuister, G.W. (2016). CcpNmr AnalysisAssign: a flexible platform for integrated NMR analysis. J Biomol NMR 66, 111–124. 10.1007/s10858-016-0060-y.

12. Maciejewski, M.W., Schuyler, A.D., Gryk, M.R., Moraru, I.I., Romero, P.R., Ulrich, E.L., Eghbalnia, H.R., Livny, M., Delaglio, F., and Hoch, J.C. (2017). NMRbox: A rsource for biomolecular NMR computation. Biophys J 112, 1529–1534. 10.1016/j.bpj.2017.03.011.

13. Pervushin, K., Riek, R., Wider, G., and Wüthrich, K. (1997). Attenuated T2 relaxation by mutual cancellation of dipole–dipole coupling and chemical shift anisotropy indicates an avenue to NMR structures of very large biological macromolecules in solution. Proc Natl Acad Sci USA 94, 12366–12371. 10.1073/pnas.94.23.12366.

14. Lescop, E., Schanda, P., and Brutscher, B. (2007). A set of BEST triple-resonance experiments for time-optimized protein resonance assignment. J Magn Reson 187, 163–169. 10.1016/j.jmr.2007.04.002.

15. Schulte-Herbrüggen, T., and Sorensen, O.W. (2000). Clean TROSY: compensation for relaxation-induced artifacts. J Magn Reson 144, 123–128. 10.1006/jmre.2000.2020.

16. Shiraishi, Y., and Shimada, I. (2023). NMR characterization of the papain-like protease from SARS-CoV-2 identifies the conformational heterogeneity in Its inhibitor-binding site. J Am Chem Soc 145, 16669–16677. 10.1021/jacs.3c04115.

17. Klemm, T., Ebert, G., Calleja, D.J., Allison, C.C., Richardson, L.W., Bernardini, J.P., Lu, B.G., Kuchel, N.W., Grohmann, C., Shibata, Y., et al. (2020). Mechanism and inhibition of the papain-like protease, PLpro, of SARS-CoV-2. EMBO J 39. 10.15252/embj.2020106275.

